# Motor guidance by long-range communication through the microtubule highway

**DOI:** 10.1101/2020.12.23.424221

**Authors:** Sithara S. Wijeratne, Shane A. Fiorenza, Radhika Subramanian, Meredith D. Betterton

## Abstract

Coupling of motor proteins within arrays drives muscle contraction, flagellar beating, chromosome segregation, and other biological processes. Current models of motor coupling invoke either direct mechanical linkage or protein crowding, which rely on short-range motor-motor interactions. In contrast, coupling mechanisms that act at longer length scales remain largely unexplored. Here we report that microtubules can physically couple motor movement in the absence of short-range interactions. The human kinesin-4 Kif4A changes the run-length and velocity of other motors on the same microtubule in the dilute binding limit, when 10-nm-sized motors are separated by microns. This effect does not depend on specific motor-motor interactions because similar changes in Kif4A motility are induced by kinesin-1 motors. A micron-scale attractive interaction potential between motors is sufficient to recreate the experimental results in a computational model. Unexpectedly, our theory suggests that long-range microtubule-mediated coupling not only affects binding kinetics but also motor mechanochemistry. Therefore, motors can sense and respond to motors bound several microns away on a microtubule. These results suggest a paradigm in which the microtubule lattice, rather than being merely a passive track, is a dynamic medium responsive to binding proteins to enable new forms of collective motor behavior.

---

Diverse cellular processes rely on coordinated activity of cytoskeletal motor proteins. For example, minifilaments made of multiple myosin motors pull actin filaments together to contract muscle.^1,2^ Similarly, dynein motors line the microtubule doublet and collectively induce the oscillatory beating of motile flagella.^3,4^ “Trains” of motors mediate intraflagellar transport, which is essential for assembly and maintenance of cilia and flagella.^5,6^ Force balance between plus- and minus-end-directed motors that crosslinks microtubules is a proposed mechanism to maintain mitotic spindle organization.^7–10^ Similarly, tug-of-war between opposite polarity motors underlies bidirectional cargo transport.^11–13^ For all of these processes, the activity of multiple motors is coupled.

Currently, the best-understood mechanisms of motor-motor coupling fall into two categories: protein crowding or mechanical linkage. Motors can be mechanically linked, either by directly binding to each other or binding to the same cargo. For example, in myosin minifilaments many motors form an ensemble that collectively generates force to contract muscles against high load.^14^ Alternatively, motors that are densely crowded on cytoskeletal filaments can have altered activity due to short-range steric interactions and/or cooperativity.^15,16^ Kinesin-1 motors form clusters due to short-range attractive interactions, for example.^17,18^ Kinesins that regulate microtubule dynamic instability typically accumulate at microtubule ends where their motility changes. For example, the activity of the kinesin-8 Kip3p is altered in dense clusters at the ends of microtubules.^19–21^ Another prototypical example is the mitotic spindle-associated kinesin-4 protein Kif4A, which forms clusters at microtubule ends (hereafter referred to as “end-tags”) and regulates microtubule length.^22–24^ Short-range interactions are well-studied and recognized as important for motor ensemble function. However, whether coupling between proteins at longer length scales contributes to the organization of motor ensembles remains unclear.

Recent work has suggested that motor and non-motor microtubule associated proteins can structurally alter the tubulin lattice. This raises the possibility of long-range coupling through the microtubule polymer. Lattice effects are proposed to influence microtubule dynamics directly or indirectly by altering the activity of regulatory proteins.^25,26^ Kinesin-1 motors have been shown to cause lattice defects^27^ and expansion^28^ and alter the binding affinity of kinesin-1 to microtubules,^29,30^ possibly due to elastic anisotropy.^31^ This effect can result in cooperative binding of kinesins to the same microtubule. Currently, whether long-distance coupling through the “medium” of the microtubule can affect motile properties of motor proteins is not known. It further remains unclear whether long-range coupling mechanisms can dynamically sense and respond to motor density on microtubules, particularly at low concentration. Finally, whether coupling between proteins at a longer length scale contributes to the formation of motor ensembles/clusters is unknown. Hence, we have a limited understanding of whether concentration-dependent long-range coupling might be a general mechanism that determines the spatial organization of motors on microtubules.

In this work, we report unexpected long-range coupling between Kif4A motors on microtubules at low density. The micron-length-scale coupling leads to a density-dependent change in Kif4A processivity and speed at picomolar motor concentration, where short-range protein-protein interactions are unlikely. The results indicate that kinesins can influence the movement of motor molecules that are widely separated on microtubules, even without physical short-range coupling, oligomerization, external binding partners, or tubulin post-translational modification. Computational modeling suggests that long range coupling is likely to affect the mechanochemical stepping cycle of the motor in addition to the binding kinetics. At higher protein concentration, motor coupling on the nanometer- and the micron-scale co-exists and results in the organization of microtubule-length-dependent Kif4A end-tags, which is not predicted for moderately processive motors like Kif4A. These observations enlarge our understanding of how the microtubule can act as a responsive medium for communication between motors separated on the micron length scale.

## Results

The kinesin-4 motor Kif4A accumulates at microtubule ends, where it binds with high affinity.^23,24^ In contrast to other highly processive kinesins or complexes, such as Kip3p^19–21^ or the PRC1-Kif4A complex,^24^ Kif4A alone is only moderately processive, with an average measured run length of about 1 *μ*m.^23^ Despite this, previously published data show that Kif4A end-tags are sensitive to overall microtubule length for microtubules up to 14 *μ*m.^24^ To understand how motors could possibly exhibit length-dependent behavior at length scales an order of magnitude larger than their average run length, we investigated the formation of end-tags by Kif4A motors.

To measure end-tag formation and its dependence on microtubule length and motor concentration, we reconstituted the activity of Kif4A on single microtubules. For these studies, we used a Total Internal Reflection Fluorescence (TIRF) microscopy assay as reported previously.^24,32^ First, rhodamine-labeled, taxol-stabilized microtubules were biotinylated and immobilized on a glass coverslip (Fig. 1A). Next, GFP-tagged Kif4A (0.02 nM) was added to the flow chamber for 5 min and then imaged. Near-simultaneous multi-wavelength imaging of rhodamine-labeled microtubules and Kif4A-GFP showed that Kif4A preferentially accumulates at the plus-end of microtubules, as observed previously (Fig. 1B).^24^ With increasing Kif4A concentration (0.02-6 nM), the length of the end-tags increases. In particular, the micron-sized end-tags at higher Kif4A concentration (4 nM) resemble those formed from the collective activity of Kif4A and PRC1 at concentrations of 1.5 nM and 0.1-0.4 nM, respectively.^24^ We measured the end-tag length and intensity over a range of filament lengths up to 13 *μ*m (Fig. 1C, S1A). The data fit well to a straight line, where the slope corresponds to the fraction of the microtubule length that is the end-tag (Figure 1D, S1B). These results show microtubule-length dependence of end-tags formed by Kif4A alone.

**Figure 1:**
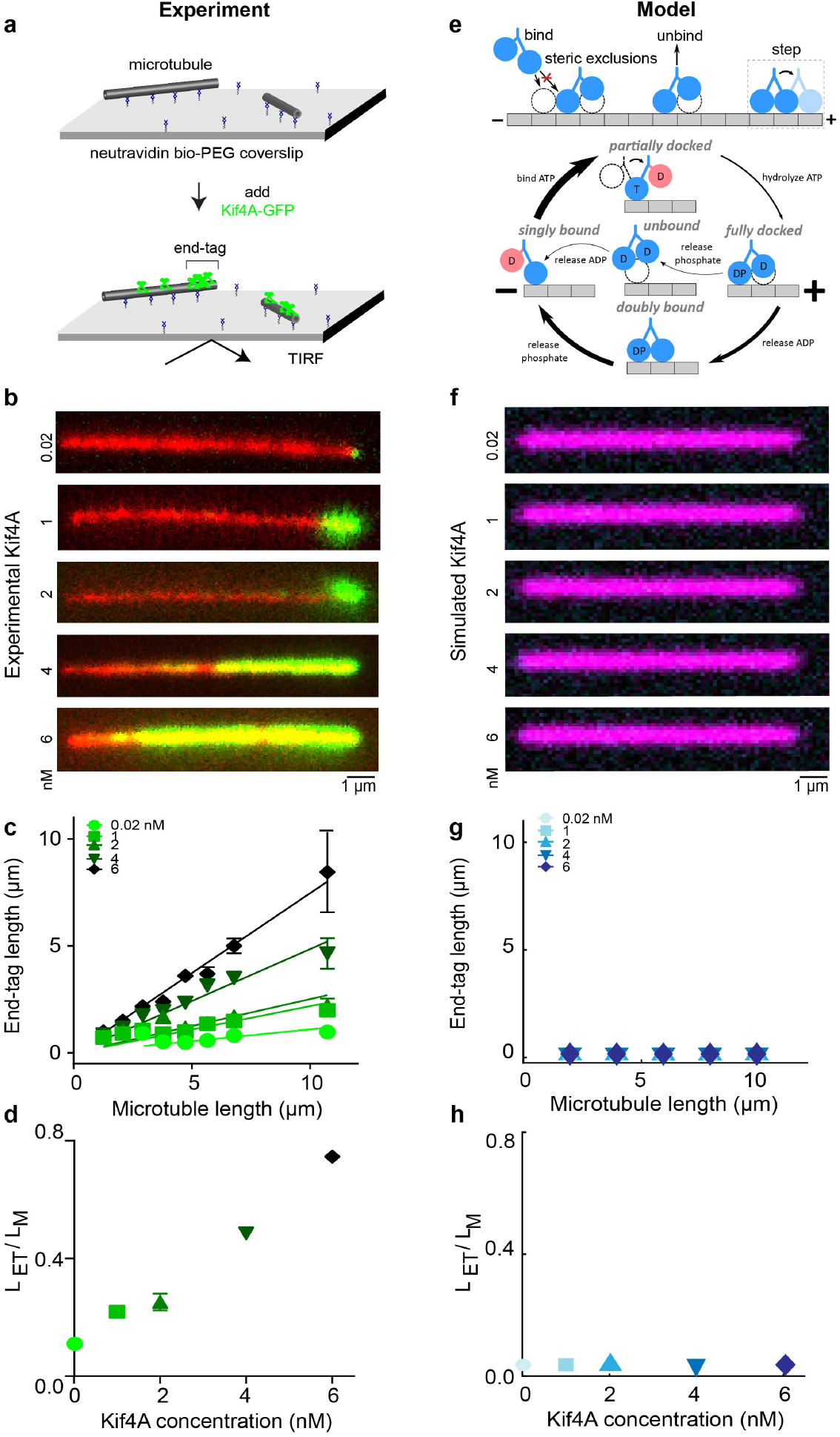
The kinesin-4 motor Kif4A forms microtubule-length-dependent end-tags, but a minimal motor model does not reproduce the experimental observations. A. Schematic of the *in vitro* assay used for examining the length dependence of Kif4A-GFP (green) on single microtubules (gray). B. Representative fluorescence micrographs showing end-tag formation with Kif4A-GFP concentration from 0.02 to 6 nM. Images show X-rhodamine labeled microtubules (red) with Kif4A-GFP (green). C. End-tag length versus microtubule length in assays with Kif4A-GFP concentration from 0.02 to 6 nM: 0.02 nM (slope 0.11 ± 0.02), 1 nM (slope 0.22 ± 0.02), 2 nM (slope 0.25 ± 0.03), 4 nM (slope 0.49 ± 0.02) and 6 nM (slope 0.75 ± 0.02). D. Slope (end-tag length divided by microtubule length) versus Kif4A concentration. E. Model overview. Motors can bind to, unbind from, and step toward the plus ends of microtubules. Steric interactions prevent more than one motor head from occupying a single site. Inset, model mechanochemical cycle. Motor heads can be bound to ADP (D), ATP (T), ADP o Pi (DP), or nothing (empty). Red coloring labels head which cannot bind to the microtubule in these states. Arrow thickness represents the relative probability of each transition. F. Simulated fluorescence images of our model using 10 micron-long microtubules and varying Kif4A concentration from 0.02 to 6 nM. G. Simulated end-tag length versus microtubule length. H. Slope (simulated end-tag length divided by microtubule length) versus Kif4A concentration.

We sought to understand how Kif4A motors with a run length of only ~1 micron can form length-dependent end-tags on microtubules that are ~10 microns long by developing a mathematical model of Kif4A motion and accumulation on microtubules (supplementary material). We developed a motor model that includes binding to and unbinding from microtubules, stepping via a mechanochemical cycle, and steric exclusion. We modeled a single protofilament of the microtubule as a one-dimensional lattice, where each 8-nm tubulin dimer is represented by a discrete binding site. This model builds on previous theory of motor accumulation on microtubules and traffic jams.^21,33–39^

To investigate how motor coupling might alter Kif4A behavior, we modeled motor stepping with a mechanochemical cycle driven by ATP hydrolysis, building on previous work.^40–46^ We constructed the model based on the kinesin-1 stepping cycle (Fig. 1E, supplementary material).^47–53^ While the details of which nucleotide state is associated with each conformational state may be different for Kif4A compared to kinesin-1, our model predictions are similar for any model that includes a basic mechanochemical cycle that facilitates asynchronous binding and unbinding of two binding heads.^49,54^ A general aspect of our model is that nucleotide hydrolysis rate determines motor velocity and the relative rates of second-head binding and first-head unbinding determine average motor processivity.

To investigate end-tag formation, we start with the premise that accumulation of end-tags requires that the motors create a crowded region at the plus-end of the microtubule. We assume that no binding site can be occupied by more than one motor head. If a motor is blocked from stepping forward by another motor in front of it, the rear head can still unbind, causing the motor to become stuck in the singly bound state. We constrained parameters of the model using motor processivity and velocity from previously published data on Xklp1,^23^ and the motor on-rate was estimated from experiments imaging the binding to and motility of Kif4A-GFP on microtubules at low motor density.^24^

In our simulations of this minimal model, end-tags do not form and motor accumulation does not vary with microtubule length (Fig. 1F-H). This is consistent with our intuition that a motor with a run length of only 1.2 *μ*m cannot show enhanced accumulation on microtubules several microns long.

The lack of end-tag formation in the model suggests that the model is missing key mechanisms, such as interactions between motors that alter their behavior in dense ensembles. We therefore examined whether cooperative interactions between motors might be required for end-tags. Previous work on kinesin-1 found that the motors cluster together on microtubules more than would be expected for purely non-interacting motors.^17,18^ These data were consistent with a short-range (nearest-neighbor) attractive interaction between motors with an estimated energy of 1.6-1.8 k_*B*_T.^17,18^ A similar short-range interaction would be expected if Kif4A can physically interact with nearby motors, perhaps by binding between motor tails. To test whether such a short-range interaction could lead to end-tags, we implemented a nearest-neighbor interaction that lowers the motor unbinding rate when motors are adjacent (supplementary material). We find that short-range cooperativity between adjacent Kif4A motors is not sufficient for end-tag formation, even if the interaction energy is increased up to 10 k_*B*_T (Fig. S2A). We also tried artificially increasing motor processivity by an order of magnitude (without short-range interactions), but end-tags still did not form (Fig. S2B). This suggests that another, unknown mechanism might alter Kif4A motility in ensembles.

To investigate how motor density on microtubules might change the motility of Kif4A, we examined the interaction of single Kif4A-GFP molecules with microtubules with varying concentration of unlabeled Kif4A (Fig. 2A). We considered two possibilities: at high protein concentration direct protein-protein interactions might alter motor processivity. Alternatively, the presence of even widely spaced motors on the microtubule might indirectly alter Kif4A processivity. To distinguish these cases, we studied low Kif4A concentration, in the picomolar range where direct protein-protein interactions are unlikely. At 20 pM, single Kif4A-GFP molecules moved only short distances before dissociation. However, the addition of small amounts of additional unlabeled Kif4A (30-400 pM) led to longer unidirectional movements of individual Kif4A-GFP molecules. When just 60 pM of unlabeled Kif4A was added, the average run length and lifetime of Kif4A increased by a factor of ~3 and ~4, respectively (Fig. 2B, C, E, F), along with a 2-fold reduction in the average velocity (Fig. 2D, G). These data suggest that even at sub-nanomolar protein concentration, where Kif4A-Kif4A interactions are not predominant, the processivity and velocity of the motor are sensitive to protein density on microtubules.

**Figure 2:**
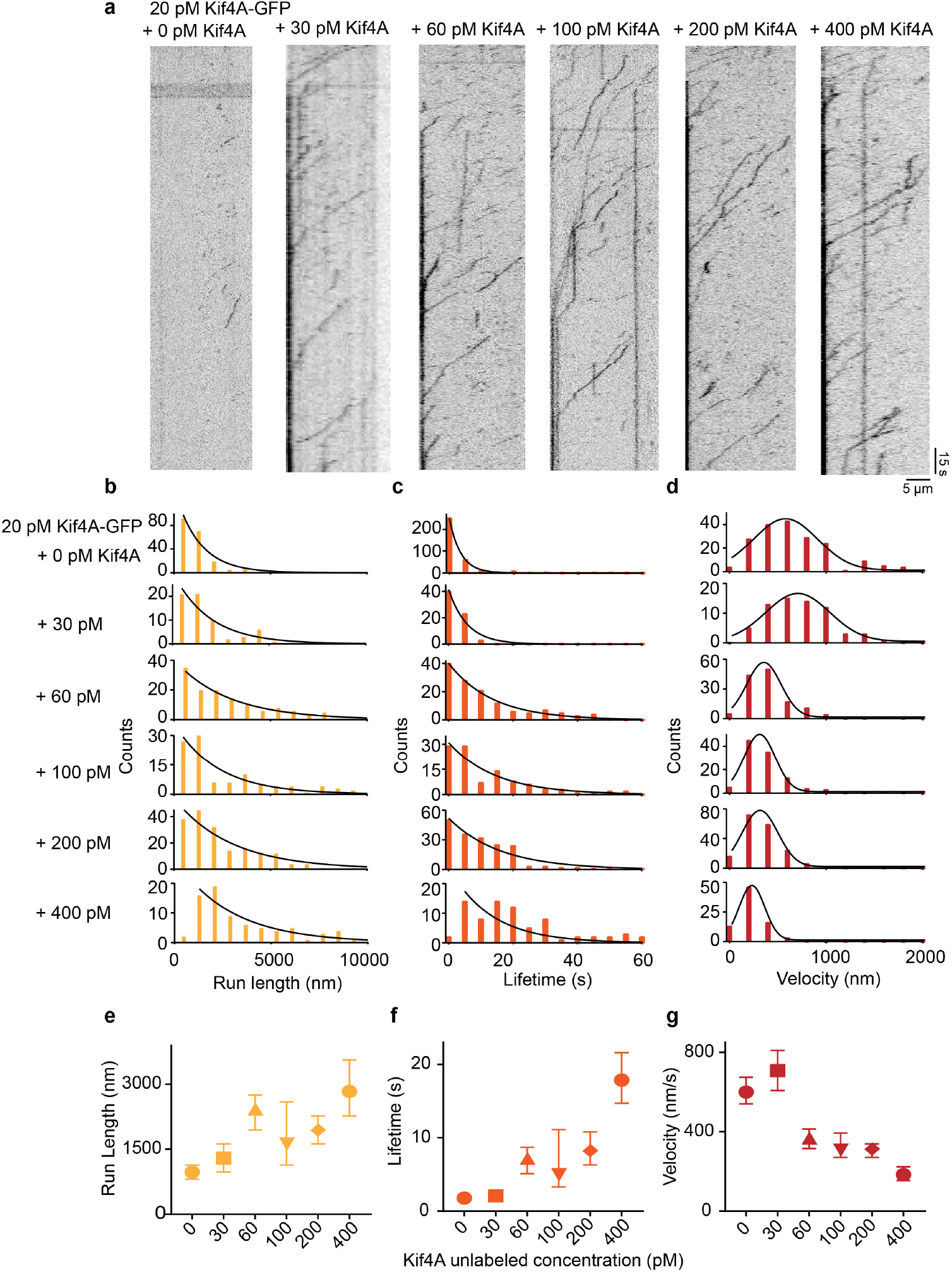
Single molecule analysis of Kif4A-GFP movement in the presence of Kif4A-unlabeled. A. Kymographs obtained from time-lapse image sequence acquired in examining microtubule interaction of Kif4A-GFP (20 pM) in presence of 0, 30, 60, 100, 200 and 400 pM Kif4A-unlabeled molecules. B-D. Histograms of the run length (B) lifetime (C) and average velocity (D) obtained from time-lapse image sequence acquired in examining microtubule interaction of Kif4A-GFP (20 pM) in presence of 0, 30, 60, 100, 200 and 400 pM Kif4A-unlabeled molecules. The run length and lifetime histograms were fit to an exponential function. The average velocity histogram was fit to a Gaussian distribution. E. Average run length versus Kif4A concentration, obtained from median in (C): 0 pM (972 nm, N=205), 30 pM (1296 nm, N=66) 60 pM (2430 nm, N=134), 100 pM (1620 nm, N=106), 200 pM (1944 nm, N=182) and 400 pM (2835 nm, N=78). F. Average lifetime versus Kif4A concentration, obtained from the median in (D): 0 pM (1.8 s, N=205), 30 pM (2.1 s, N=66), 60 pM (7.2 s, N=134), 100 pM (4.9 s, N=106), 200 pM (8.3 s, N=182) and 400 pM (17.9 s, N=78). G. Average velocity versus Kif4A concentration, obtained from the median in (E): 0 pM (599 nm/s, N=205), 30 pM (707 nm/s, N=66), 60 pM (368 nm/s, N=134), 100 pM (306 nm/s, N=106), 200 pM (313 nm/s, N=182) and 400 pM (183 nm/s, N=78). The error bars represent 95% confidence interval of the median.

We then used our mathematical model to ask what mechanisms of motor interaction might explain the surprising result that Kif4A run length, lifetime, and speed vary with density at picomolar concentration. We began with the short-range attractive potential discussed above (Fig. S2A). As expected, short-range cooperativity alone is not sufficient to reproduce the low-density data (Fig. 3A-D), even if the interaction strength is increased to 10 k_*B*_T (Fig. S3).

**Figure 3:**
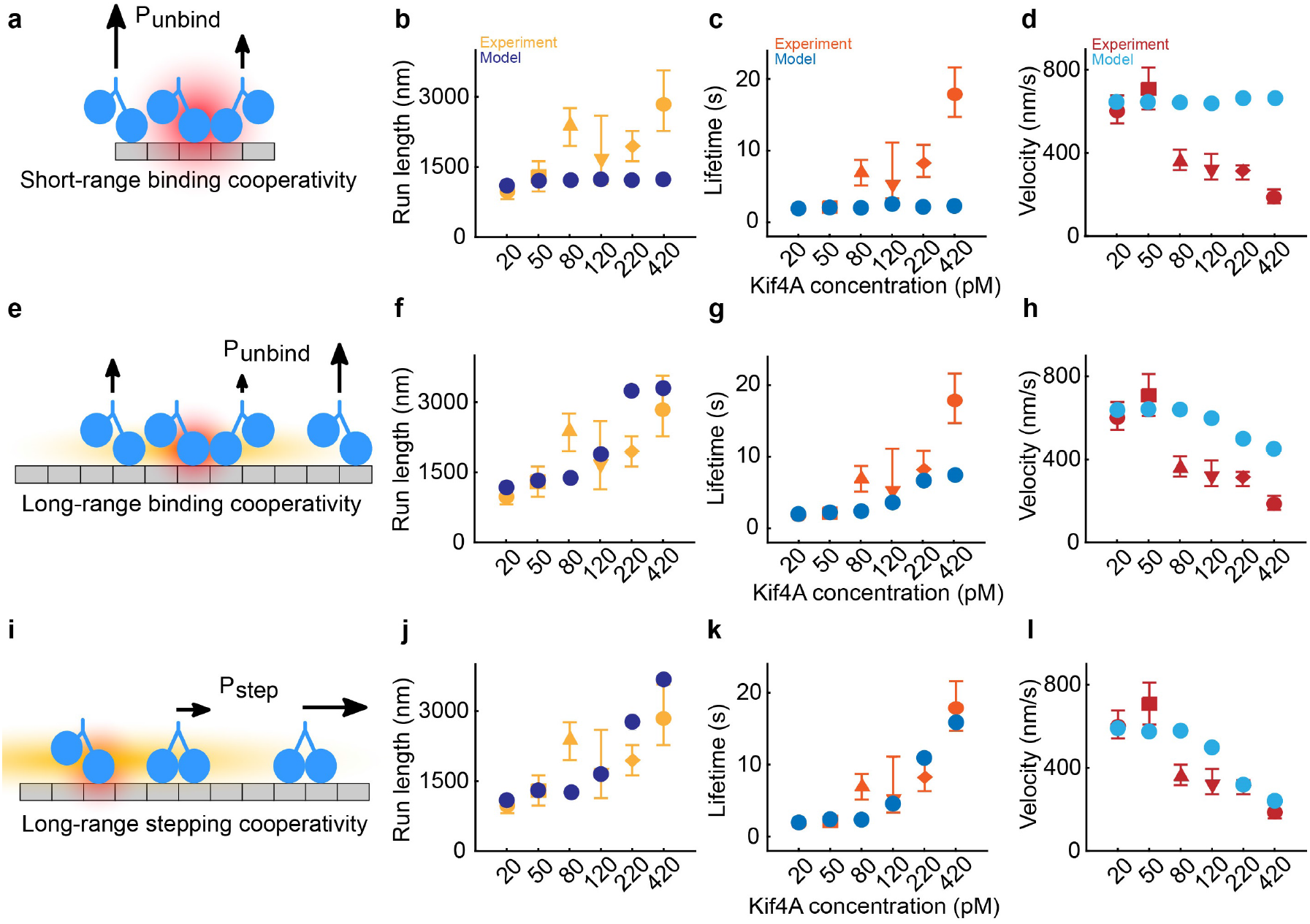
Long-range cooperativity is essential in the model and must affect motor stepping to fully reproduce the experimental results. A. Schematic of short-range cooperativity. The red cloud shows the range of the interaction, and the length of arrows shows relative event probability. In the model, short-range interactions decrease motor unbinding but does not affect binding. B-D. Motor run length, lifetime, and velocity versus motor concentration for simulation (blue) and experiment (orange, red). E. Schematic of long-range cooperativity. The orange cloud shows the range of the interaction (not to scale, simulated range is 8 *μ*m). Long-range cooperativity affects motor binding and unbinding and is implemented in addition to short-range cooperativity. F-H. Motor run length, lifetime, and velocity versus motor concentration for simulation (blue) and experiment (orange, red). I. Schematic of long-range cooperativity in stepping. The long-range cooperativity lowers motor velocity in addition to binding and unbinding. Short-range cooperativity remains in the model. J-L. Motor run length, lifetime, and velocity versus motor concentration for simulation (blue) and experiment (orange, red).

Therefore, we considered the possibility of long-range interactions between motors. Previous work found enhanced binding of kinesin-1 motors near other motors, an effect with a remarkably long range of 6 *μ*m.^29^ Such a long-range interaction could change motor-microtubule binding kinetics at the low density of our experiments, suggesting it as a possible mechanism of motor coupling. We modeled long-range motor interactions by adding an attractive quadratic potential between motors with a range of several microns (supplementary material). We assumed that the interaction potential from multiple motors is additive up to a saturation energy around 5 k_*B*_T. To mimic the interaction observed for kinesin-1, we first implemented an effect on motor binding kinetics: the long-range potential increases the binding rate and decreases the unbinding rate of other motors (Fig. 3E-H). We note that this effect was included in addition to short-range cooperativity discussed above. After fitting the three long-range interaction parameters (potential strength, range, and saturation energy) to the experimental data, we found that long-range cooperativity in the model predicts changes in motor motility at low density qualitatively similar to those found experimentally. However, the best-fit model did not show strong quantitative agreement with the data, suggesting that long-range interactions that alter motor-microtubule binding kinetics partially but not fully explain the data. Therefore, we considered whether additional mechanisms might better explain our low-density results.

The data show that Kif4A speed slows by a factor of 2–3 as motor density is increased. This is surprising in the absence of dense traffic jams where steric effects predominate, suggesting the possibility that a long-range interaction between motors might alter motor mechanochemistry. To be consistent with our data, such a mechanism would lead to decreased speed of motors coupled through the long-range interaction. Many mechanisms of slowing motor stepping are possible. One possibility would be changes in the doubly bound motor off- and ATP-binding rates due to the long-range interaction (Figure 1E, supplementary material). With this addition, the best-fit model agrees well with our experimental data (Fig 3I-L). Other mechanisms for which the long-range interaction causes motors to slow their stepping will have similar effects. These results show that long-range coupling between motors that affects both binding kinetics and motor stepping can explain the alteration in Kif4A motility with density.

If the long-range motor coupling between Kif4A molecules is mediated by changes to the microtubule lattice, it would be predicted to occur with change in density of other kinesins besides Kif4A. We measured the motility of single Kif4A-GFP molecules in the presence of increasing concentration (30-400 pM) of unlabeled *D. melanogaster* kinesin-1 dimers (amino acids 1-401; referred to as K401). K401 is a minimal kinesin-1 dimer comprising the motor domain, neck linker and the first dimerization coiled-coil domain.^55^ When the total motor concentration was increased by adding unlabeled K401, single Kif4A-GFP molecules moved more processively and exhibited long unidirectional runs (Fig. 4). When we added 60 pM of unlabeled K401 to an assay with 20 pM Kif4A-GFP, the average run length and lifetime increased by a factor of 2 and 4, respectively, while the average velocity decreased. These results show Kif4A processivity and velocity are sensitive to the density of a non-interacting motor of a different kinesin family. Because K401 lacks the C-terminal cargo binding domains typically responsible for protein-protein interactions in kinesins, these data support the idea that the motor coupling interactions that impact Kif4A motility likely do not arise from short-range protein-protein interactions. Additionally, single-molecule intensity measurements confirm that oligomerization or similar phenomena are not occurring (Fig. S4). Together, our experimental results with varying Kif4A and K401 density and our modeling results suggest that motility of single Kif4A motors is modulated by coupling of motors separated by microns along the microtubule.

**Figure 4:**
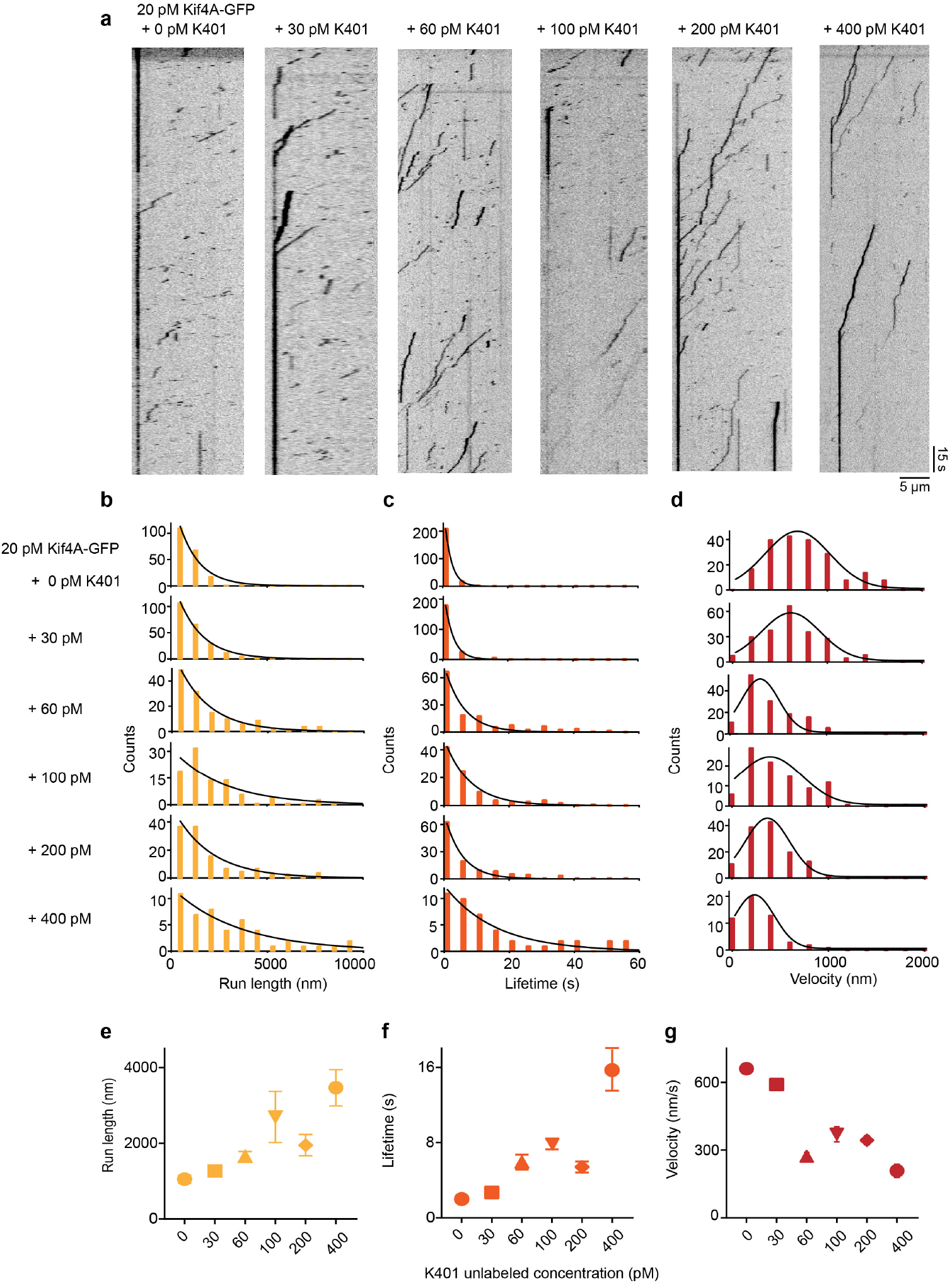
Single-molecule analysis of Kif4A-GFP movement in the presence of K401-unlabeled. A. Kymographs obtained from time-lapse image sequence of microtubules with Kif4A-GFP (20 pM) in presence of 0 pM K401-unlabeled, 60 pM K401-unlabeled, 100 pM K401-unlabeled, 200 pM K401-unlabeled and 400 pM K401-unlabeled. B-D. Histograms of the run length (B), lifetime (C), and average velocity (D) obtained from time-lapse image sequence of Kif4A-GFP (20 pM) in presence of 0 pM K401-unlabeled, 60 pM K401-unlabeled, 100 pM K401-unlabeled, 200 pM K401-unlabeled and 400 pM K401-unlabeled. Run length and lifetime histograms were fit to an exponential function. The average velocity histogram was fit to a Gaussian distribution. E. Average run length versus K401 concentration, obtained from the median in (B): 0 pM (810 nm, N=202), 30 pM (972 nm, N=228), 60 pM (1458 nm, N=140), 100 pM (1620 nm, N=96), 200 pM (1296 nm, N=129) and 400 pM (2430 nm, N=51). F. Average lifetime versus K401 concentration, obtained from the median in (C): 0 pM (1.2 s, N=202), 30 pM (1.5 s, N=228), 60 pM (4.2 s, N=140), 100 pM (3.9 s, N=96), 200 pM (3.6 s, N=129) and 400 pM (9 s, N=51). G. Average velocity versus K401 concentration, obtained from the median in (D): 0 pM (706 nm/s, N=202), 30 pM (583 nm/s, N=228), 60 pM (324 nm/s, N=140), 100 pM (402 nm/s, N=96), 200 pM (360 nm/s, N=129) and 400 pM (233, N=51). The error bars represent 95% confidence interval of the median.

Comparison of the low-density experimental results to our model suggests that a combination of short- and long-range cooperativity can explain the increase in Kif4A processivity and decrease in velocity as motor density increases on microtubules. We next asked whether these interactions are sufficient to explain the formation of end-tags on microtubules at high density. To do this we used the model to predict high-density Kif4A behavior with no free parameters: we increased the Kif4A concentration in the simulation, while maintaining the model parameters determined by fitting the low-density data. Remarkably, the model predicts end-tags that quantitatively match those found experimentally (Fig. 5A-D). The model end-tag length increases both with microtubule length and motor concentration, as in experiments. To further dissect which interactions in the model are most important for end-tag formation, we turned off parts of the model individually. Removing individual cooperative interactions from the model (corresponding to turning off short-range cooperativity, long-range cooperativity that affects binding, or long-range cooperativity that affects mechanochemistry) decreases end-tag formation (Fig. S5). This suggests that the combination of both long- and short-range motor coupling that we identified in the low-density model combine to allow Kif4A to form end-tags. In the model, the long-range interaction helps increase processivity so that motors reach the end of the microtubule, while the short-range interaction prevents unbinding to maintain motors in the end-tag.

**Figure 5:**
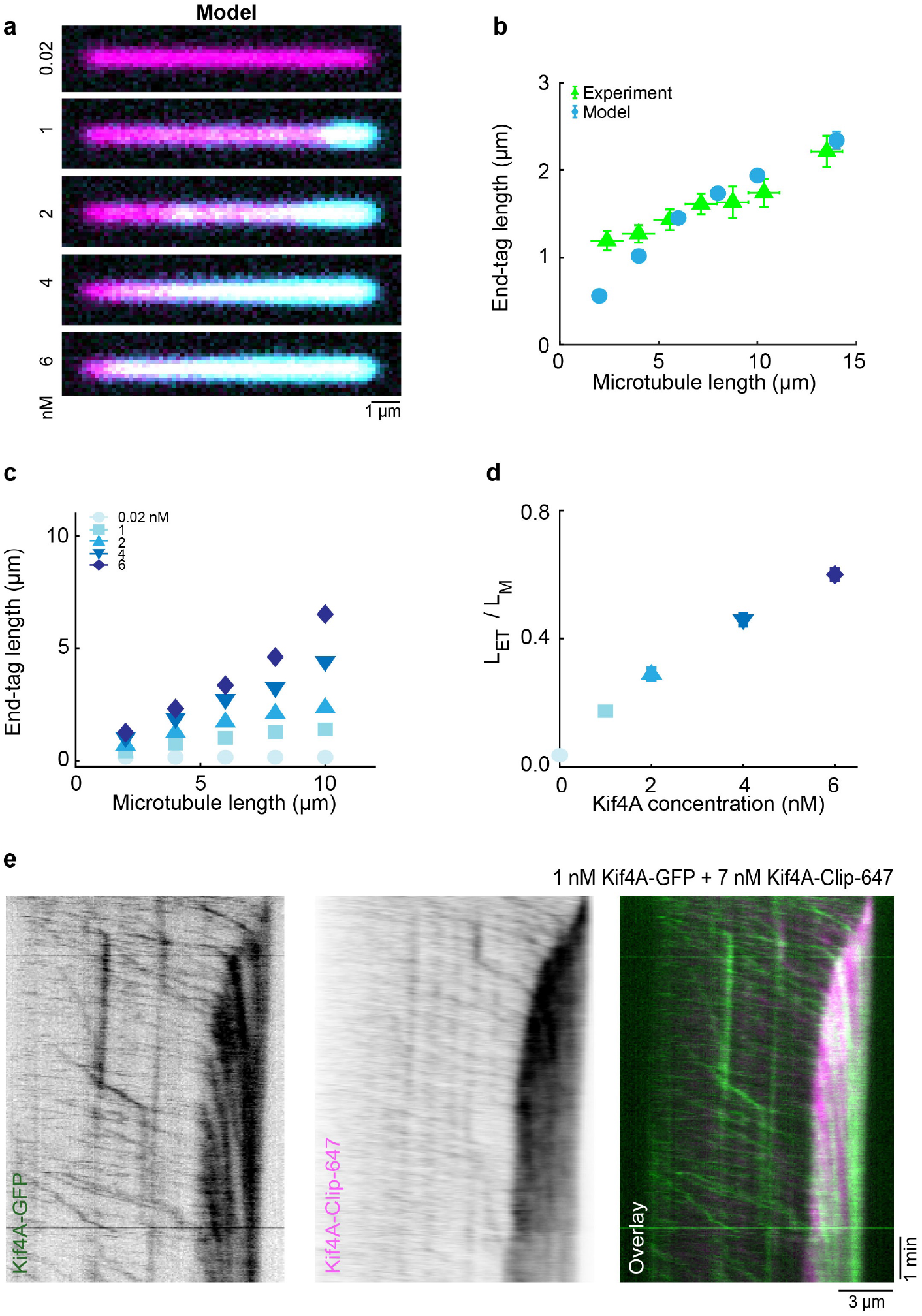
The computational model with long-range cooperativity that fits low-density experiments predicts length-dependent end-tags and Kif4A motility changes with no free parameters. A. Simulated fluorescence images created from the model of 10 *μ*m microtubules with varying Kif4A concentration. B. End-tag length versus microtubule length from simulations with long-range cooperativity (blue) and experiment (green). C. Simulated end-tag length versus microtubule length with long-range cooperativity. D. Simulated slope of end-tag length divided by microtubule length versus Kif4A concentration. E. Kymographs obtained from time-lapse sequence acquired in spiking experiments of Kif4A-Clip-647 (7 nM) in presence of Kif4A-GFP (1 nM) on single microtubule.

Based on our modeling results, we propose that long-range motor coupling between Kif4A molecules that increases processivity and lowers velocity contributes to the formation of dense end-tags on microtubules. The model predicts that near and in the end-tag, the bound lifetime of Kif4A increases and its speed drops. To examine whether these changes occur in end-tags, we directly visualized processive movement of single Kif4A molecules at high protein concentration by spiking in Kif4A-GFP (1 nM) with Kif4A-Alexa647 (7 nM) while observing end-tag formation in real time (Fig. 5E). In these experiments, end-tag formation initiates at the microtubule plus-end and grows toward the minus-end until a steady-state end-tag length is established. Outside the end-tag, motors move processively with long plus-end-directed runs (~5000 nm or longer). Motor velocity in the untagged region of the microtubule was 110 nm/s, but upon encountering the high-density end-tag, Kif4A-GFP slowed to 25 nm/s. These results suggest that, consistent with our model predictions, end-tag formation occurs through an increase in Kif4A processivity at high concentration along with a reduction in velocity and dissociation in the end-tags.

## Discussion

Here we describe motor communication that spans microns without the usual physically linked assembly. We discovered these interactions for Kif4A, a kinesin-4 motor known to cluster at microtubule ends, but the changes in Kif4A motility can also be induced by kinesin-1 motors. Our findings suggest that long-range coupling of motors through the microtubule can impact both the binding and mechanochemistry of motor proteins at low density. Thus, the microtubule can act as an allosteric medium for microtubule-associated kinesins to sense the number of motors on the same microtubule. This coupling can set up a positive feedback loop whereby motors adaptively increase their processivity, even at picomolar concentration where motors are typically microns apart.

Our work expands the way we think about microtubules as a medium for allosteric coupling and how it can affect motor proteins. Long-range coupling between motors is required to explain motor-density-dependent changes in processivity at picomolar concentration. At these concentrations, a minimal computational model that utilizes a conventional-kinesin stepping cycle and a short-range attractive potential (such as that arising from traffic jams at high concentration and protein-protein interaction) cannot explain the experimental observations. This is because the widely spaced motors hardly ever come close enough to each other for short-range interactions to occur (Movie S1). Long-range interactions may have the advantage of allowing motors with individually low processivity to change their collective behavior without any hindrance resulting from direct physical interactions. Interestingly, the increase in processivity from long-range interactions at low density leads to clustering of motors at the microtubule ends at higher concentration, resulting in the formation of microtubule-length-dependent end-tags. While motor binding proteins can increase motor processivity and lead to clustering, long-range coupling provides a motor-autonomous mechanism to increase protein density on the microtubule.

The observed effects on motor velocity as a function of density at low concentration cannot be fully explained by a mechanism whereby the long-distance coupling affects only the binding-unbinding kinetics of the motor-microtubule interaction. By contrast, our computational model suggests that long-range cooperativity affects motor mechanochemistry directly. This model satisfactorily reproduces the experimental data at both low (0.002 nM) and high (6 nM) Kif4A concentration with no change in model parameters, which argues that both single-motor properties (such as processivity) and emergent behavior of motor ensembles (such as end-tag formation) require long-range coupling between motors. This differs from previous results on kinesin-1 both because of the strong effect on Kif4A motility and because the changes occur for much lower motor concentration for Kif4A (tens of picomolar) versus kinesin-1 (tens of nanomolar).^29,30^ As a result, long-range motor coupling can drive end-tag formation for Kif4A but not kinesin-1. Thus, the long-range coupling mechanism can increase the diversity in the outcome of the collective motor activity on microtubules depending on the properties of individual proteins.

Previous work has proposed that conformational changes in tubulin heterodimers mediated by the binding of microtubule-associated proteins can act as an allosteric coupler within the microtubule lattice.^25,26^ Our findings broaden the scenario in which these effects are relevant by suggesting that the molecular and structural alterations mediating microtubule allosteric coupling do not require a saturated microtubule lattice and are reversible on the time scale of seconds. Therefore, long-distance coupling can be achieved without requiring long-term alteration of the microtubule lattice. In contrast, mechanisms such as tubulin isoform diversity, post-translational modification, and protofilament register shifts are long-lived or irreversible structural/biochemical changes to the microtubule. Transient motor-autonomous long-distance coupling might confer a unique advantage, as microtubules can quickly respond to changes in protein concentration to regulate kinesin motility.

The long-range coupling we describe has significant implications for motor-based cellular processes because only a small number of motors need to bind on a microtubule to trigger a cascade (Movie S2). For example, in the context of intracellular transport, long-range coupling may facilitate changes in velocity, motor force-velocity relation, or the outcome of tug-of-war between opposing motors, on a specific subset of cellular microtubules. Beyond coupling between motors, the long-range effects may also impact microtubule ends to control dynamic instability. For example, increased Kif4A processivity can increase the protein concentration at microtubule ends, which could then alter the polymerization of dynamic microtubules.^23^ Thus low-density, long-distance interactions may allow motors to self-organize without physical short-range coupling, oligomerization, binding partners, or tubulin post-translational modifications. This kind of coupling can make kinesin motors more adaptable, allowing them to perform different functions depending on the surrounding environment and local motor concentration in cells.

**Figure 6:**
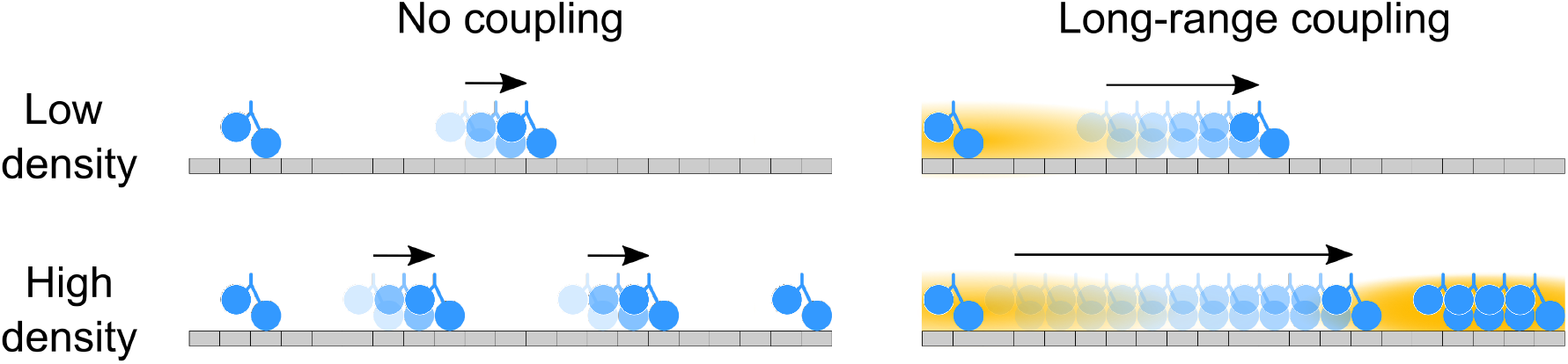
Illustration of effects of long-range motor coupling. Schematic shows motors (blue) moving on microtubule (gray) with interaction region (orange cloud, not to scale). Length of arrows represents motor run length (not to scale). (Top left) Non-interacting motors do not affect the run length or velocity of other motors. (Top right) Long-range interactions mean that Kif4A changes the run-length and velocity of widely separated motors on the same microtubule. Our theory suggests that this long-range coupling affects not only binding kinetics, but also motor mechanochemistry. (Lower left) Non-interacting motors do not change their motility or collective behavior at higher density. (Lower right) Long-range coupling promotes the formation of Kif4A end-tags at high density. The microtubule therefore responds dynamically to motor binding and alters the behavior of other motors, allowing new forms of collective motor behavior.

Our results suggest a new view in which the microtubule is not a passive highway on which motors move, but instead a responsive medium that couples motors moving along it. Motors moving along microtubules may therefore be analogous to other physical systems in which new forms of collective behavior occur due to coupling through a medium, such as Cooper-paired electrons in a superconductor,^56^ diffusion of atoms of the surface of a crystal,^57^ liquid-liquid phase separation in an elastic gel,^58,59^ and interactions of active particles through a granular medium.^60^

## Methods

### Protein purification

Recombinant proteins (Kif4A, Kif4A-GFP, Kif4A-CLIP and K401) were expressed and purified as described previously.^24, 61^ For CLIP protein labeling, purified protein was incubated with CLIP-Surface™ 647 in a 1:3 (protein:dye) molar ratio, at 30°C for 1 hr, followed by incubation at 4°C overnight. Unbound dye was removed by repeated dilution and centrifugation through an Amicon^®^ Ultra-15 Centrifugal Filter Unit (Millipore Sigma), prior to size exclusion chromatography. A comparison of the absorbance of pure labeled protein at 650 nm and 280 nm yielded a labeling efficiency of 15-20%.

### Microtubule polymerization

Taxol-stabilized rhodamine-labeled microtubules were prepared with biotin tubulin as described previously.^24^ Briefly, GMPCPP seeds were prepared from a mixture of unlabeled bovine tubulin, X-rhodamine-tubulin and biotin tubulin, which were diluted in BRB80 buffer (80 mM PIPES pH 6.8, 1.5 mM MgCl_2_, 0.5 mM EGTA, pH 6.8) and mixed together by tapping gently. The tube was transferred to a 37°C heating block and covered with foil to reduce light exposure. The biotinylated microtubules were incubated for 1 hr 45 min. Afterwards, 100 μL of warm BRB80 buffer was added to the microtubules and spun at 75000 rpm, 10 min, and 37°C to remove free unpolymerized tubulin. Following the centrifugation step, the supernatant was discarded, and the pellet was washed by round of centrifugation with 100 μL BRB80 supplemented with 20 μM taxol. The pellet was resuspended in 16 μL of BRB80 containing 20 μM taxol and stored at room temperature covered in foil.

### In vitro fluorescence microscopy assay

The microscope slides (Gold Seal Cover Glass, 24 × 60 mm, thickness No.1.5) and coverslips (Gold Seal Cover Glass, 18 × 18 mm, thickness No.1.5) were cleaned and functionalized with biotinylated PEG and non-biotinylated PEG, respectively, to prevent nonspecific surface sticking, according to standard protocols. Flow chambers were built by applying two strips of double-sided tape to a slide and applying to the coverslip. Sample chamber volumes were approximately 6-8 *μ*L.

Experiments were performed as described previously.^24^ To visualize the accumulation of Kif4A on microtubules, rhodamine-labeled biotinylated were immobilized in a flow chamber coated with neutravidin (0.2 mg/ml). Next, Kif4A-GFP and and 1 mM ATP were flushed into the flow chamber in assay buffer (BRB80 buffer supplemented with 1 mM TCEP, 0.2 mg/ml k-casein, 20 *μ*M taxol, 40 mg/ml glucose oxidase, 35 mg/ml glucose catalase, 0.5% *β*-mercaptoethanol, 5% sucrose and 1 mM ATP). The flow cell was incubated for 10 min before taking snapshots of the microtubule and GFP channel. To visualize single molecules, Kif4A-GFP and 1 mM ATP were flowed into the chamber in assay buffer and a time-lapse sequence of images was immediately acquired at a rate of 0.3 frames/s. Data were collected for 2-4 min. Experiments with K401 were performed using the same method.

All experiments were performed on Nikon Ti-E inverted microscope with a Ti-ND6-PFS perfect focus system equipped with an APO TIRF 100x oil/1.49 DIC objective (Nikon). The microscope was outfitted with a Nikon-encoded x-y motorized stage and a piezo z-stage, an sCMOS camera (Andor Zyla 4.2), and two-color TIRF imaging optics (Lasers: 488 nm and 561 nm; Filters: Dual Band 488/561 TIRF exciter).

### Image analysis

ImageJ (NIH) was used to process the image files. Briefly, raw time-lapse images were converted to tiff files. A rolling ball radius background subtraction of 50 pixels was applied to distinguish the features in the images more clearly. From these images, individual microtubule single molecule events were picked and converted to kymographs by the MultipleOverlay and MultipleKymograph plug-ins (J. Reitdorf and A. Seitz). We then extracted parameters such as run length and lifetime, and calculated the average velocity (run length/lifetime), for each single molecule track. We only included moving single molecule events and excluded stalled events from the analysis.

## Acknowledgements

We thank Matthew Glaser, Dick McIntosh, Michael Shelley, and Daniel Needleman for useful discussions. We thank Nandini Mani for help with protein purification. This work was funded by NSF grants DMR-1725065 (MDB), NIH grants R01GM124371 (MDB) and 1DP2GM126894 (RS). Simulations used the Summit supercomputer, supported by NSF grants ACI-1532235 and ACI-1532236.

