## Supplemental text and figures for "Motor guidance by long-range communication through the microtubule highway"

**This PDF file includes:**

Supplementary text
Supplementary references
Table S1
Figures S1 to S5
Captions for Movies S1 and S2

**Other supplementary materials for this manuscript include the following:**

Movies S1 and S2

### Computational Model

**Microtubules.** We model a single protofilament of the microtubule, and each 8-nm tubulin dimer corresponds to a discrete binding site on a 1-D lattice. All binding sites have the same base binding affinity (before adding effects of interactions). Microtubules are polar and have sites that correspond to a plus- and minus-end. The microtubule is assumed fixed in space and hydrodynamic interactions between motors are neglected.

**Motors.** Each motor consists of two independently binding heads connected by a rigid linkage. Motors are implicit in solution and randomly drawn from a reservoir upon binding. Motors bind to the microtubule with rate $k_{on}cn_u$, where $n_u$ is the number of unoccupied binding sites available and $c$ is the bulk concentration of motors. The first motor head to bind is always leading, i.e., closer to the plus-end than the unbound head. Motor heads can be bound to ADP, ATP, ADP•Pi, or no ligand (empty). While in solution, both motor heads are assumed to be bound to ADP. Upon binding to the microtubule, the first motor head releases ADP and becomes empty. While empty or ATP-bound, motor heads are strongly bound and cannot unbind from the microtubule. ATP binding to the empty head occurs at rate $k_{ATP}$ and induces a conformational change that swings the second (unbound) head forward, partially docking the motor. The first (bound) head then hydrolyzes ATP to ADP•Pi at rate $k_{hydro}$, which fully docks the motor. Next, the motor can either unbind its first (bound) head at rate $k_{off,\,i}$, which terminates its run, or bind its second (unbound) head to the microtubule at rate $k_{on}c_{eff}$, making it doubly bound and continuing its run. The ratio between $k_{off,\,i}$ and $k_{on}c_{eff}$ determines how many steps occur on average before the motor unbinds. While the motor is doubly bound, the rear head unbinds with rate $k_{off,ii}$ and the front head cannot unbind. Upon rear head unbinding, the motor is again singly bound state, after having advanced forward one site along the microtubule.

**Steric exclusion.** No two motor heads can occupy the same binding site at the same time. We neglect any steric effects off the microtubule lattice. For example, we assume the trailing head of a singly-bound motor can always swing forward during a conformational change, regardless of how crowded the surrounding solution is.

**End-pausing.** We model end-pausing by disabling the partial-docking conformational change induced by ATP binding when a motor head is bound to the microtubule plus-end. The trailing head can still re-bind to the microtubule as long as the site remains unoccupied.

**Effects of binding energy change on kinetics.** Changes in binding and unbinding rates due to motor interaction energy follow Boltzmann statistics. If a state change such as binding or unbinding changes the energy by $\Delta U$, the equilibrium distribution of the states correctly samples the Boltzmann distribution if the association (or forward) rate $k_a$ and dissociation (or reverse) rate $k_d$ are altered as $k_a = k_a^0 exp[-\beta(1-\lambda)\Delta U]$ and $k_d = k_d^0 exp[\beta\lambda\Delta U]$, where $\beta = 1/k_B T$ and the dimensionless parameter $\lambda$ typically lies in the range $0 \le \lambda \le 1$. For an attractive interaction between motors, the free energy change decreases upon binding, making $\Delta U$ negative. This means the association rate increases and the dissociation rate decreases relative to no interaction. The parameter λ weights how the interaction affects binding and unbinding. If $\lambda = 1$ unbinding is strongly affected by the interaction while binding is not affected; in the other limit $\lambda = 0$, binding is strongly affected by the interaction and unbinding is not affected. When $\lambda = 1/2$, binding and unbinding are equally affected by the interaction. Note that the dissociation constant $K_D = k_d/k_a$ is $K_D = K_D^0 exp[2\beta\Delta U]$ with the interaction, consistent with the expectation that the equilibrium is independent of $\lambda$.

**Short-range binding cooperativity.** Short-range binding cooperativity in our model is a neighbor-neighbor interaction with a range of one lattice site. The attractive potential has a negative energy of magnitude $\epsilon$ (written in units of $k_B T$). We set $\lambda = 1$ so that binding is unaffected by the interaction, giving $k_a = k_a^0$ and $k_d = k_d^0 exp[-n\varepsilon]$, where $n = 0, 1, 2$ is the number of neighbors of the motor head. We assume that motor heads of the same motor do not interact cooperatively, so doubly-bound motor heads can have a maximum of one neighbor. Only bound motor heads lead to an interaction.

**Long-range binding cooperativity.** To model long-range binding cooperativity, we use the interaction potential $E(x) = \frac{1}{2}\alpha x^2 - E_0$, which is negative (attractive) up to the cutoff distance $D$ where $E(D) = 0$, so the interaction does not extend beyond $D$. Here $x$ is the distance in nm along the 1-D lattice, $E_0$ is the strength of the interaction energy in units of $k_B T$, and $\alpha$ is an effective spring constant set by $E(D) = 0$. The effects of multiple motors superpose up a maximum $E^*$, which represents the upper limit at which long-range interactions saturate. We choose $\lambda = 1/2$ so that both binding and unbinding are equally affected, $k_a = k_a^0 exp[-\frac{1}{2}\Sigma E_i]$ and $k_d = k_d^0 exp[\frac{1}{2}\Sigma E_i]$, where $\Sigma E_i$ is the sum of all motor energies in units of $k_B T$ up to the maximum $E^*$. Unless otherwise stated, this long-range binding cooperativity is always implemented alongside the short-range binding cooperativity described above.

**Long-range stepping cooperativity.** Motor interactions could occur via multiple mechanisms, which would have similar effects as long as the interaction lowers the motor stepping speed. We modeled one plausible hypothesis for the mechanism: the internal necklinker tension that couples motor heads could be decreased by the interaction. To implement this, we extended the model of necklinker tension during motor stepping of Andreasson et al. (Ref. 1). Our changes modify the rear-head unbinding rate of doubly bound motors to $k_{off,ii} = k_{off,ii}^0 exp[\beta F_i \sigma_{off}]$ and introduce a new doubly bound → partially docked pathway with rate $k_{ATP,ii} = k_{ATP}^0 exp[-F_i \sigma_{ATP}]$, where $F_i$ is the internal necklinker tension, and $\sigma_{off}$ and $\sigma_{ATP}$ are unbinding and ATP-binding distance parameters, respectively. If ATP binds to the front head, the rear head detaches from the microtubule and swings forward, skipping the singly bound state. The values estimated for kinesin-1 by Andreasson et al. are $F_i = 26$ pN, $\sigma_{off} = 0.35$ nm, and $\sigma_{ATP} = 4.6$ nm. Using $k_{off,ii} = 260$ s$^{-1}$ and $k_{ATP,ii} = 5000$ s$^{-1}$ also reported by Andreasson et al. leads to doubly bound off- and ATP-binding rates of $2375$ and $1.18 \times 10^{-9}$ per second, respectively. Normally, this force-dependence would not significantly alter our simulations, since the internal force is already taken into account in our doubly bound off-rate and the doubly bound ATP binding rate is nearly zero. However, in our model the long-range binding cooperativity above changes the internal necklinker tension. We implement this by multiplying the doubly bound ATP binding and off-rate by the $exp[-\frac{1}{2}\Sigma E_i]$ and $exp[\frac{1}{2}\Sigma E_i]$, respectively.

**Data fitting.** Model parameters are shown in Table S1. We fit model parameters to the low-density Kif4A data using nonlinear least-squares optimization from the SciPy python library, using a trust region reflective algorithm. The fit gave an initial estimate of the six parameters $\varepsilon$, $D$, $E_0$, $E^*$, $\sigma_{off}$, and $\sigma_{ATP}$. Final parameter adjustment by hand gave the final parameter set.

**Approach to steady state.** Simulations run for a time $t_e$ to pre-equilibrate (approach steady state in binding). The fractional occupancy of the microtubule, defined as the number of occupied binding sites divided by the total number of binding sites, is averaged. At the end of pre-equilibration, fractional occupancy is calculated and stored each timestep. Every $t_c$ seconds after, the average and standard deviation of fractional occupancy are calculated. If the change in average fractional occupancy is less than the standard deviations added in quadrature, the system is considered equilibrated. At this point, data collection begins and continues until the end of the simulation.

**Kinetic Monte Carlo algorithm.** We implement the model in a hybrid tau-skipping kinetic Monte Carlo (kMC) simulation, which samples from the binomial and Poisson distributions to predict the number of events that will occur each fixed timestep. Events in each timestep are executed in a random order on randomly selected members of appropriate populations. Since multiple events can affect a single population, e.g., binding of the second head and unbinding of the first head for singly bound motors, we ensure that no two events can act on the same motor. Our simulation executes the following algorithm each timestep:

1. Scan over all active motors and sort into appropriate populations
2. Sample appropriate statistical distribution for each possible event. If the probability of the event is constant, such as for example ATP binding, use the binomial distribution. If the probability varies each timestep, such as for example motor unbinding in the presence of long-range cooperativity, sum the total probabilities and use the Poisson distribution as described in Ref. 2.
3. If two or more events target the same motor, iteratively sample a random number to discard at random until only one remains. Each event probability is used as a relative weight when sampling to discard.
4. Shuffle event order and execute each event on particles randomly sampled from the appropriate population.
5. Increment simulation time by $dt$.

Data collection begins after a simulation reaches steady state.

**End-tag length analysis.** Microtubule occupancy data are averaged over all points to determine steady-state fractional occupancy. This occupancy data are smoothed with moving average window 250-nm wide, chosen to match the diffraction limit of visible light. Starting from the plus-end of the microtubule, the first site past peak occupancy with a slope less than half the peak slope is considered the end-tag boundary.

**Simulated fluorescence images.** The motor fractional occupancy is averaged for 0.5 seconds, to mimic image collection time. We then apply a Gaussian blur filter to the average motor occupancy to create the cyan channel of the image. We repeat this process for a fully occupied array to represent tubulin as the magenta channel. The yellow channel is made up of random noise. The simulated fluorescence image is a merge of these three channels, and movies are image sequences.

**Table S1.** Model parameter values, based on final fit to the data as shown in Fig. 3C. In the simulations shown in fig 3A and 3B, $F_i = 0$, doubly bound off-rate $k_{off.ii}$ = 2375 $s^{-1}$ (Ref. 1). When modeling K401, the hydrolysis rate is increased to 135 $s^{-1}$ to increase the average velocity at 20 pM concentration to 800 nm/s.

| Quantity | Parameter | Value | Notes |
|---|---|---|---|
| **General** | | | |
| $k_B T$ | Thermal energy | 4.1 pN·nm | Room temperature |
| $t$ | Total simulation time | $35 - 170$ min | Longer time for lower Kif4A concentration |
| $t_e$ | Pre-equilibration time | 100 s | |
| $t_c$ | Equilibration check interval | 10 s | |
| $dt$ | Timestep | $2 \times 10^{-5}$ s | |
| $\Delta$ | Site size | 8 nm | Length of a tubulin dimer |
| $L$ | Microtubule length | $2 - 14$ $\mu$m | Experimental values |
| **Motors** | | | |
| $c$ | Bulk motor concentration | $0.002 - 6.0$ nM | Experimental values |
| $c_{eff}$ | Effective concentration of second head while singly bound | $4 \times 10^6$ nM | Assumes the second head can explore the volume of a quarter-sphere with a necklinker 7.5 nm long |
| $k_{on}$ | Per-site motor binding rate | $3.6 \times 10^{-4}$ nM$^{-1}$s$^{-1}$ | Estimated from 16 kymographs of Kif4A at 20 pM; assumes 6 of the 13 protofilaments can bind motors |
| $k_{ATP}$ | ATP binding rate | 5000 $s^{-1}$ | Andreasson et al. *eLife* (2015) |
| $k_{hydro}$ | ATP hydrolysis rate | 95 $s^{-1}$ | Chosen to give motors an average velocity of 600 nm/s |
| $k_{off,i}$ | Singly bound unbinding rate | 8 $s^{-1}$ | Chosen to give motors an average processivity of 1.2 $\mu$m |
| $k_{off,ii}$ | Doubly bound unbinding rate | 260 $s^{-1}$ | Andreasson et al. *eLife* (2015) |
| **Motors - binding cooperativity** | | | |
| $\epsilon$ | Neighbor-neighbor energy | 1.6 $k_B T$ | Best fit strength of short-range attractive potential |
| $D$ | Quadratic potential cutoff | 8 $\mu$m | Best fit range of long-range attractive potential |
| $E_0$ | Quadratic potential strength | 0.95 $k_B T$ | Best fit amplitude of long-range potential |
| $E^*$ | Quadratic potential ceiling | 4.75 $k_B T$ | Best fit maximum energy of long-range potential from multiple motors |
| **Motors - stepping cooperativity** | | | |
| $F_i$ | Internal necklinker tension | 26 pN | Andreasson et al. *eLife* (2015) |
| $\sigma_{off}$ | Distance parameter for rear head unbinding | 0.125 nm | Best fit; compare to estimate of 0.35 nm for kinesin-1 from Andreasson et al. *eLife* (2015) |
| $\sigma_{ATP}$ | Distance parameter for front head ATP binding | 1.15 nm | Best fit; compare to estimate of 4.6 nm for kinesin-1 from Andreasson et al. *eLife* (2015) |

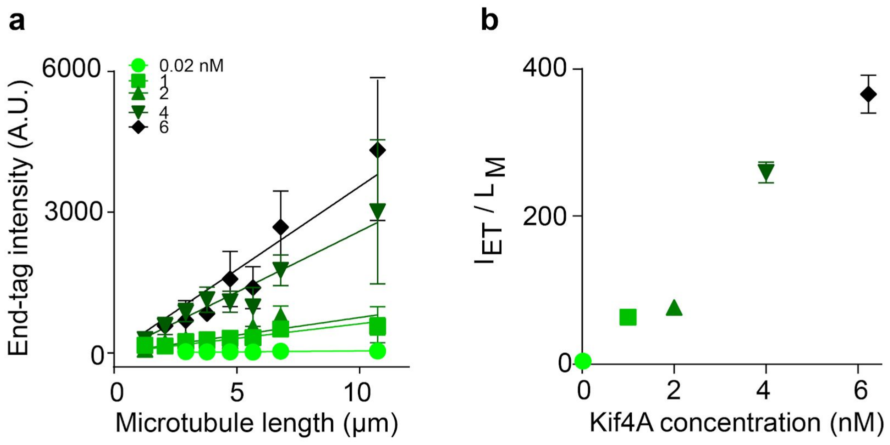


**Supplementary Figure 1.** (A) End-tag intensity versus microtubule length from assays with Kif4A-GFP concentration from 0.02 to 6 nM: 0.02 nM (circles: slope 4.44 ± 0.50), 1 nM (squares: slope 62.2 ± 4.39), 2 nM (triangles: slope 75.6 ± 9.39), 4 nM (inverted triangles: slope 259 ± 14.2) and 6 nM (diamonds: slope 355 ± 26.0). (B) Slope of plots of end-tag intensity versus microtubule length from (A) as a function of Kif4A concentration.

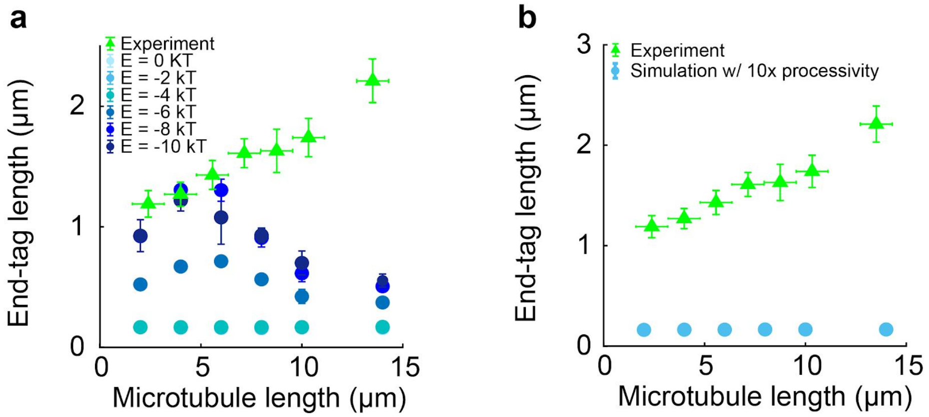


**Supplementary Figure 2.** (A) End-tag length versus microtubule length from simulations with short-range cooperativity of varying strength (blue circles) and experiment (green triangles). (B) End-tag length versus microtubule length from simulations with motor processivity increased by a factor of 10 (blue circles) and experiment (green triangles).

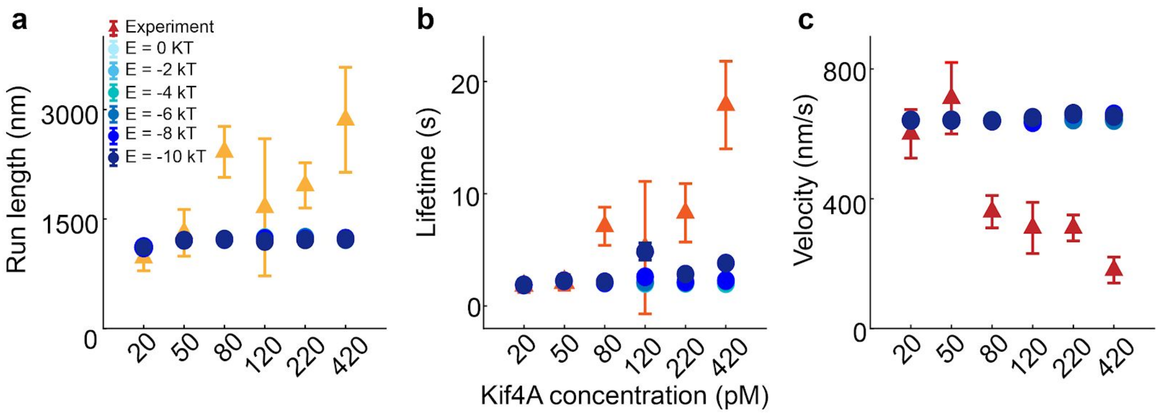


**Supplementary Figure 3. (**A) Run length, (B) lifetime and (C) velocity versus Kif4A concentration from simulations with varying energy of short-range interaction (blue) and experiment (orange, red).

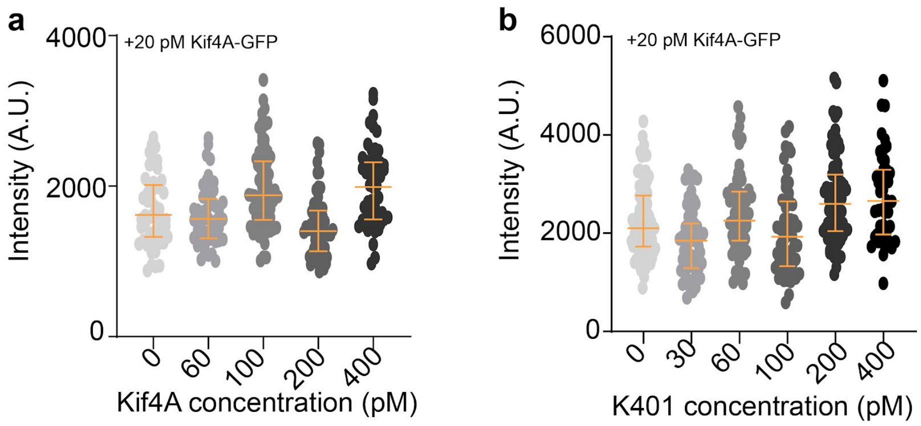


**Supplementary Figure 4.** (A) Distribution of single molecule GFP intensity for experiments with unlabeled Kif4A concentration 0-400 pM in the presence of 20 pM Kif4A-GFP. (B) Distribution of single molecule GFP intensity for experiments with unlabeled K401 concentrations 0-400 pM in the presence of 20 pM Kif4A-GFP.

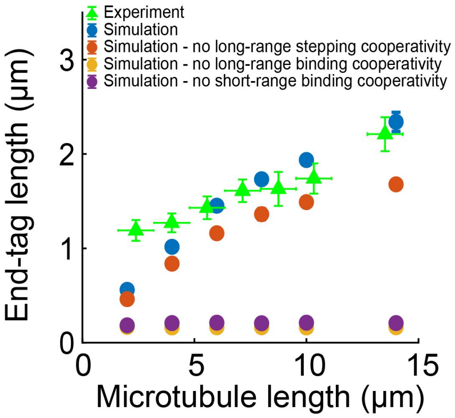


**Supplementary Figure 5.** End-tag length versus microtubule length from simulations with no short-range binding cooperativity (purple), no long-range binding cooperativity (yellow), no long-range stepping cooperativity (orange), no long range stepping cooperativity (blue) and experiment (green).

**Movie S1 (separate file).** Simulated fluorescence microscopy movie with 80 pM Kif4A. The brightness of individual motors is chosen to make individual motors clearly visible.

**Movie S2 (separate file).** Simulated fluorescence microscopy movie with 120 pM Kif4A. The brightness of individual motors is chosen to make individual motors clearly visible. A cascade of binding events occurs around t = 375 seconds. This type of cascade is not seen in ~10,000 seconds of the 80 pM simulation of Movie S1.
